# Computational investigation of Scutellarein derivatives as an inhibitor against triple-negative breast cancer by Quantum calculation, and drug-designed approaches

**DOI:** 10.1101/2023.03.06.531447

**Authors:** Shopnil Akash, Farjana Islam Aovi, Md. A.K Azad, Ajoy Kumer, Unesco Chakma, Md Rezaul Islam, Nobendu Mukerjee, Md. Mominur Rahman, Imren Bayıl, Summya Rashid, Rohit Sharma

**Affiliations:** Faculty of Allied health Science, Department of Pharmacy, Daffodil International University, Dhaka-1207, Bangladesh; Laboratory of Computational Research for Drug Design and Material Science, Department of Chemistry, European University of Bangladesh, Dhaka-1216, Bangladesh; Department of Electrical and Electronics Engineering, European University of Bangladesh, Gabtoli, Dhaka-1216, Bangladesh; Department of Microbiology; West Bengal State University, West Bengal, Kolkata-700126, India; Department of Health Sciences, Novel Global Community Educational Foundation, Hebersham, NSW, Australia; Department of bioinformatics and computational biology, Gaziantep University, Turkey; Department of Pharmacology and Toxicology, College of Pharmacy, Prince Sattam Bin Abdulaziz University, P.O. Box 173, Al-Kharj 11942, Saudi Arabia; Department of Rasa Shastra and Bhaishajya Kalpana, Faculty of Ayurveda, Institute of Medical Science, Banaras Hindu University, Varanasi-221005, India

**Keywords:** Metastasis, Tumors, Molecular docking, Molecular dynamics, DFT, ADMET, triple-negative breast cancer

## Abstract

Triple-negative breast cancer accounts for 10-15% of all breast tumors (TNBC). Triple-negative breast cancer lacks either estrogen or progesterone receptors (ER or PR), producing either too little or too much HER2. (All three tests result in “negative” results for the cells.) These cancers are more common in women under 40 who are Black or have the BRCA1 mutation. TNBC differs from other types of invasive breast cancer in that it has fewer treatment options, a worse prognosis, and grows and spreads more quickly (outcome). So, first of all, the protein of triple-negative breast cancer was collected from the PDB database having the most stable configuration, and a natural bioactive molecule, Scutellarein, was selected. Scutellarein is well-known to possess anti-cancer properties, so its derivatives were chosen to design anticancer drugs through computational tools. In this case, the functional group has applied and modified structural activity relationship methods. Then, the pass prediction score was taken, which indicates the probability of active (Pa) and the probability of inactive (Pi). After that, other *in-silco* approaches, such as the ADMET parameter, and quantum calculation by Density Functional Theory (DFT), have been conducted. Finally, molecular docking and dynamics have been evaluated against TNBC to determine the binding affinity and stability. Scutellarein derivatives (DM03 at -10.7 kcal/mol, DM04 at -11.0 kcal/mol) have been reported to have a maximum tendency for binding against TNBC. Besides, the molecular dynamic simulation was performed at 100ns and described by root-mean-square deviation (RMSD) and root-mean-square fluctuation (RMSF), which are much more stable compounds. The pharmacokinetics specifications for a suitable therapeutic candidate were satisfied by these molecules, like as non-carcinogenic, minimal to aquatic and non-aquatic toxicity. Almost all the molecules are highly soluble in the aqueous system. These all-computational data suggested that they might be suitable as a medication for the inhibition of TNBC, and further experimental studies should be carried out.

## 1.0 Introduction

Breast cancer is a difficult challenge for the global public health community[1] and the disease’s growing prevalence. It seems to substantially affect the lives of a large number of females globally by breast cancer[2, 3]. Among different types of breast cancer, triple-negative breast cancer (TNBC), one of the breast cancer subtypes with negative expression of progesterone (PR), estrogen (ER), and human epidermal growth factor receptor-2 (HER2), spreads quickly to other body regions which has a higher chance of early decline and death than other subtypes of cancer[4].

Approximately one million women are diagnosed with breast cancer each year, and TNBC accounts for approximately 15–20 percent [5, 6]. According to epidemiological research findings, TNBC is most often diagnosed in young premenopausal women under the age of 40. These make up around 15–20% of all breast cancer patients[7]. The survival duration for people diagnosed with TNBC is lower than that of patients diagnosed with other breast cancer subtypes. Forty percent of fatalities are during the first five years following identification[8, 9]. TNBC is resistant to endocrine therapy and other types of molecularly therapeutic strategies due to the distinct genetic phenotype that makes it susceptible to these treatments[10-12]. As a result, chemotherapy is the primary therapeutic option; nevertheless, the effectiveness of traditional chemoradiotherapy is relatively poor[10]. In certain countries, bevacizumab has been administered with chemotherapeutic medications to treat TNBC; however, patients did not experience a substantial improvement in their overall survival time due to this treatment[13, 14]. Treatment choices for triple-negative breast cancer are more limited than those for other kinds of invasive breast cancer. This is due to the fact that cancer cells do not possess the estrogen or progesterone receptors or enough of the HER2 protein for hormone treatment or targeted HER2 medicines to be effective against the disease[15-17]. So, even though TNBC has been more invasive in women and has affected numerous women worldwide, there is no presently acknowledged treatment for targeting this TNBC [18, 19].

The chemical compound known as scutellarin is classified as a flavone, which belongs to the phenolic family, and it has a history of use in innovative medicine containing this compound. It has been demonstrated that scutellarin may stimulate different types of cancer cells to undergo apoptosis when tested in vitro. Additionally, scutellarin has been shown to have a neuroprotective effect on nerve cells, which are negatively impacted by estrogen[20]. Scutellarin was also found to have a strong inhibitory effect on a drug-resistant strain of HIV-1 cell-to-cell multiplication when tested in a laboratory[21], as well as vigorous antibacterial and antifungal activity[22]. Table displayed efficiency of scutellarin on the proliferation of several cancer cell lines.

So, the perspective of this research is to develop and identify a potential drug candidate for the treatment of TNBC by structural modification of scutellarin. In this case, the most powerful computational method is applied, and performed different types of investigation to validate and establish scutellarin as a potential medication against TNBC. The main advantage of computational investigation, it may reduce the time, cost, and resources. These techniques may also lower the chances of failure during clinical or pre-clinical trials[29].

Scutellarin is the glycosyloxyflavone which has a significant role as an antineoplastic agent. It binds with progesterone (PR), estrogen (ER) receptor, and impedes the development of cancer cells, which the body finally eliminates. In Fig.1, TNBC cell development occurs, and gradually spread, but, when, scutellarin bind with the estrogen receptor, the production of abnormal cell growth is stopes, the production of healthy cell is started.

**Figure 1.**
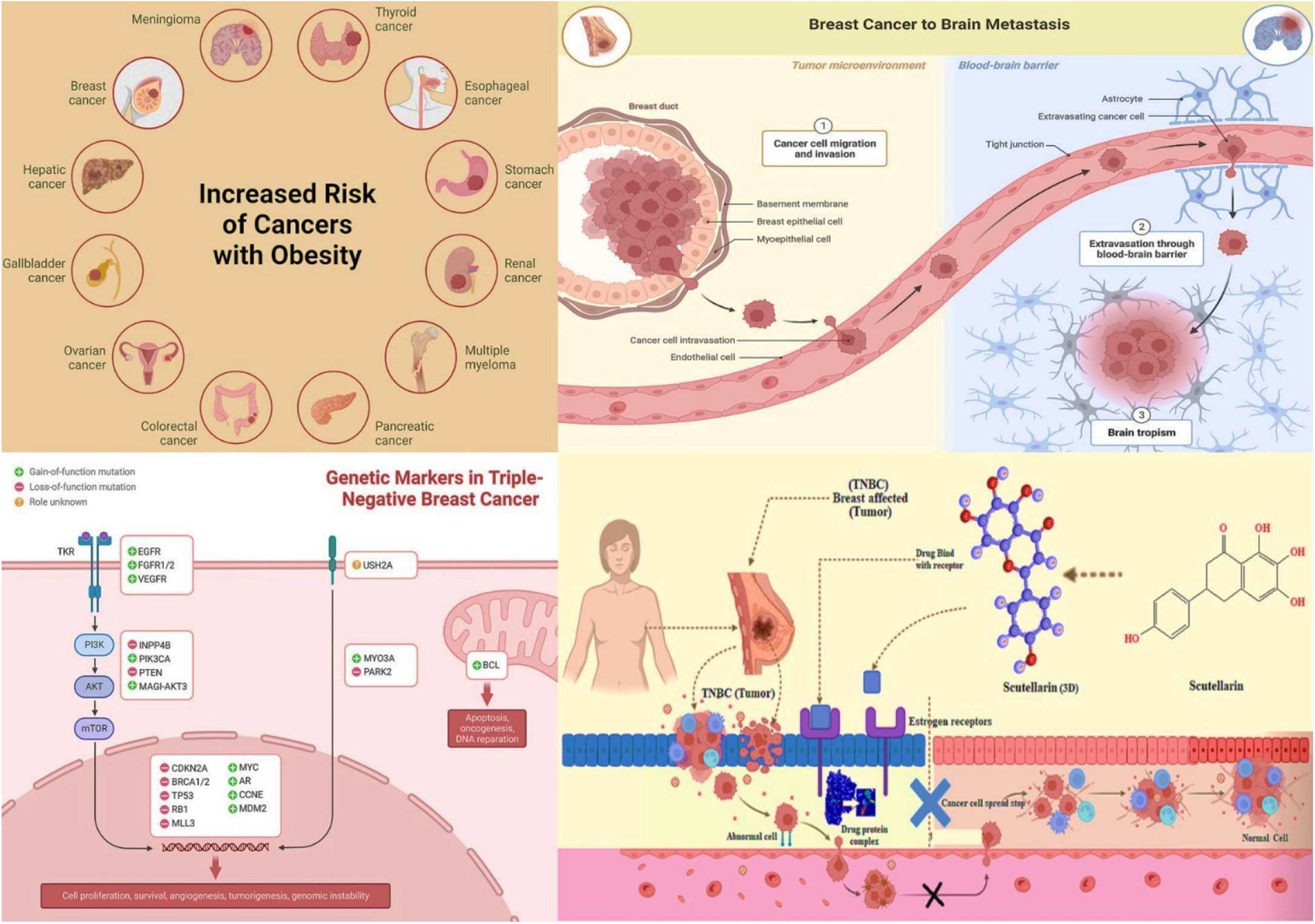
Risk factor, genetic markers, and probable mechanism of TNBC (Created with Biorender.com).

## 2. Computational method and working procedure

### 2.1. Ligand Preparation and optimization

For the preparation of ligands. Firstly, prepare the structure through Hyperchem 6.03 software[30], optimize crystal structures through material studio 08, and implement the DFT (density functional theory) from the DMol3 code[31-33]. The B3LYP functional and 6-31G++ basis sets were implemented in DMol3 code to get such exact outcomes attributed to the electronegative atom oxygen in the systems[34]. These analytical tools were implemented to establish the frontier molecular orbitals (HOMO, LUMO) and the amplitude of the HOMO and LUMO following optimization. The optimized molecules were exported as PDB file types in order to be used for subsequent computer work, such as molecular docking, molecular dynamic, and ADMET. The magnitude of chemical reactivity and characteristics are approximated by applying relevant and approved algorithms, and the results are listed in Table 4. such as energy gap, Δε = εLUMO – εHOMO; ionization potential, I = −εHOMO; electron affinity, A = −εLUMO; electronegativity, χ = (I + A)/2; chemical potential, μ = − (I + A)/2; hardness, η = (I - A)/2; electrophilicity, ω = μ2 /2η; softness, S = 1/η. The following equation is applied for the calculation of chemical descriptors.

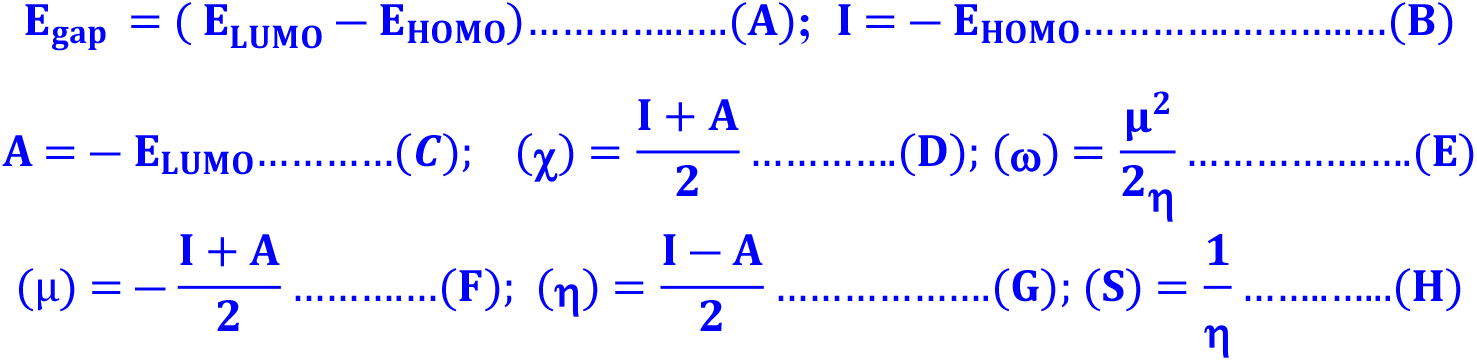

### 2.2 Lipinski Rule and Pharmacokinetics

Swiss ADME online database has been implemented to evaluate the specified compounds’ Lipinski Rule and Pharmacokinetics before the molecular docking experiment(http://www.swissadme.ch) [31]. Lipinski Rule and Pharmacokinetics may be characterized as a complicated balance of several chemical characteristics and structural factors that indicate whether a newly synthesized molecule is close to existing medications. These features are marked by hydrophobicity, drug likeness, hydrogen bonding characteristics, molecule size and molecular weight, bioavailability, and many more.

### 2.3 Determination of ADMET Profile

There is a higher prevalence of failure in developing new drugs due to inadequate pharmacokinetic and safety features. Computational techniques may be able to assist in reducing these concerns. Regarding pharmacokinetic characteristics, pkCSM could be an alternative approach to predicting ADMET features. This is the most preferable resource for estimating ADMET (absorption, distribution, metabolism, excretion, and toxicity) parameters. The report and research documented that ADMET is beneficial for predicting the pharmacokinetics of biomolecules before going to clinical or pre-clinical trials [35, 36]. So. using this online system pkCSM “https://biosig.lab.uq.edu.au/pkcsm/” online tool, the ADMET feature was evaluated and analyzed[37].

### 2.4 Method for Molecular Docking

For docking analysis, the initial three-dimensional (3D) structural triple-negative breast cancer responsible protein (PDB ID: 7L1X & PDB ID 5HA9) was collected from the protein data bank. After that, the Pymol program version V2.3 (https://pymol.org/2/) was implemented to purify the proteins such as water molecules, and unexpected ligands or heteroatoms were removed and obtained a fresh protein. Then, they were saved as PDB files[38] and wrapped the molecules with different size of grid box such as center, X = 10.2287, Y = -0.1393, Z = 15.5457, dimension (Å), X = 65.598, Y = 53.702, Z = 54.163 for protein (PDB ID 7L1Z and PDB ID 5HA9). After that, the proteins were imported into the PyRx application and converted into macromolecules; similarly, all the ligands were imported and converted in pdbqt format. Finally, molecular docking has been performed through AutoDock Vina[39]. Then, the dock file was imported into the Pymol application to generate a complex structure. The docked complexes were transferred into Discovery Studio version 2020 for further analysis and visualization of the active amino acid residues and proteins ligands pocket diagram [40]. The three-dimensional structure of the targeted protein is displayed in **Fig 2**.

**Figure 2.**
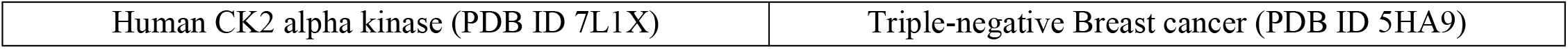
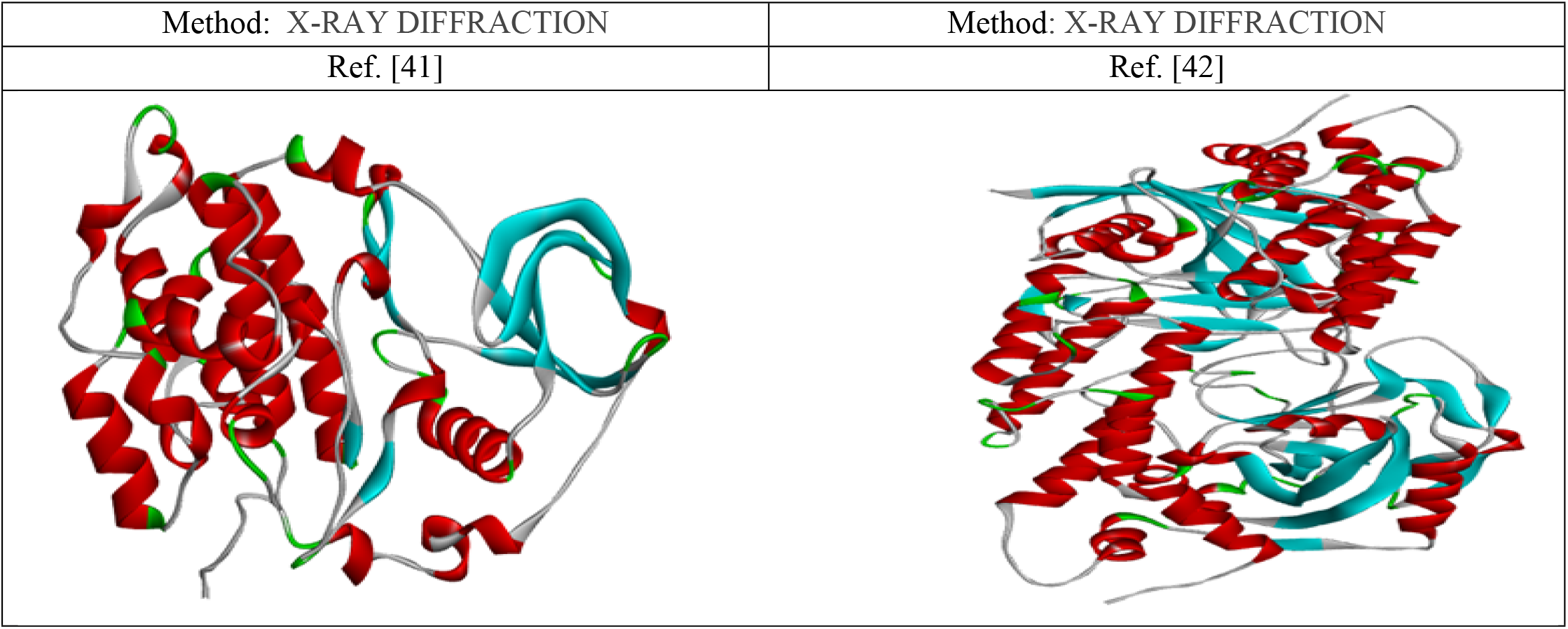
Three-dimensional protein structure of triple-negative breast cancer and its basic features.

### 2.5 Molecular Dynamic simulations (MD)

MD simulations’ purpose is to determine chemical compounds’ stability in different physiological conditions. So, the highly configured desktop computers equipped with NAMD software version 2.11 were implemented and performed in batch mode or real-time with a live view to run the MD simulation up to 100 ns [43-45]. The AMBER14 force field was implemented in the holo-form (drug-protein) of the MD simulation to provide the optimum fitting or binding pose and stabilization of the ligand-protein docking[46]. The complete system was adjusted using 0.9 percent NaCl at 298 K temperature and adding a water solvent. During the simulation, a cubic cell was transmitted within 20 Å on either side of the system and the corresponding boundary conditions. After simulation, the root means square deviation (RMSD) and root mean square fluctuation (RMSF) were analyzed using the visual molecular dynamics (VMD) software[47].

## 3. Result and Discussions

### 3.1 Designing derivatives of scutellarin

The primary compound was scutellarin which has previously been documented as anti-cancer, antimicrobial, and many more pharmacological effects. So, based on the previous investigation and literature review. The Scutellarein structure has been modified to predict the potential anti-cancer efficiency against targeted triple-negative breast cancer. Since the structure-activity relationship is a well-known approach for developing and designing new and innovative molecules to assist in the discovery or synthesis of a novel drug to get desired qualities and activities. The different position of the functional group of Scutellarein has been substituted by the Benzene ring, OCH_3,_ & NH-CH_2_-CH_2_-OH to obtain better efficacy and potential drug for inhibiting triple-negative breast cancer. The main objective is to identify how different functional groups impact pharmacological activity on Scutellarein against triple-negative breast cancer. The modified chemical structures of triple-negative breast cancer are shown in **Fig.3**.

**Figure 3.**
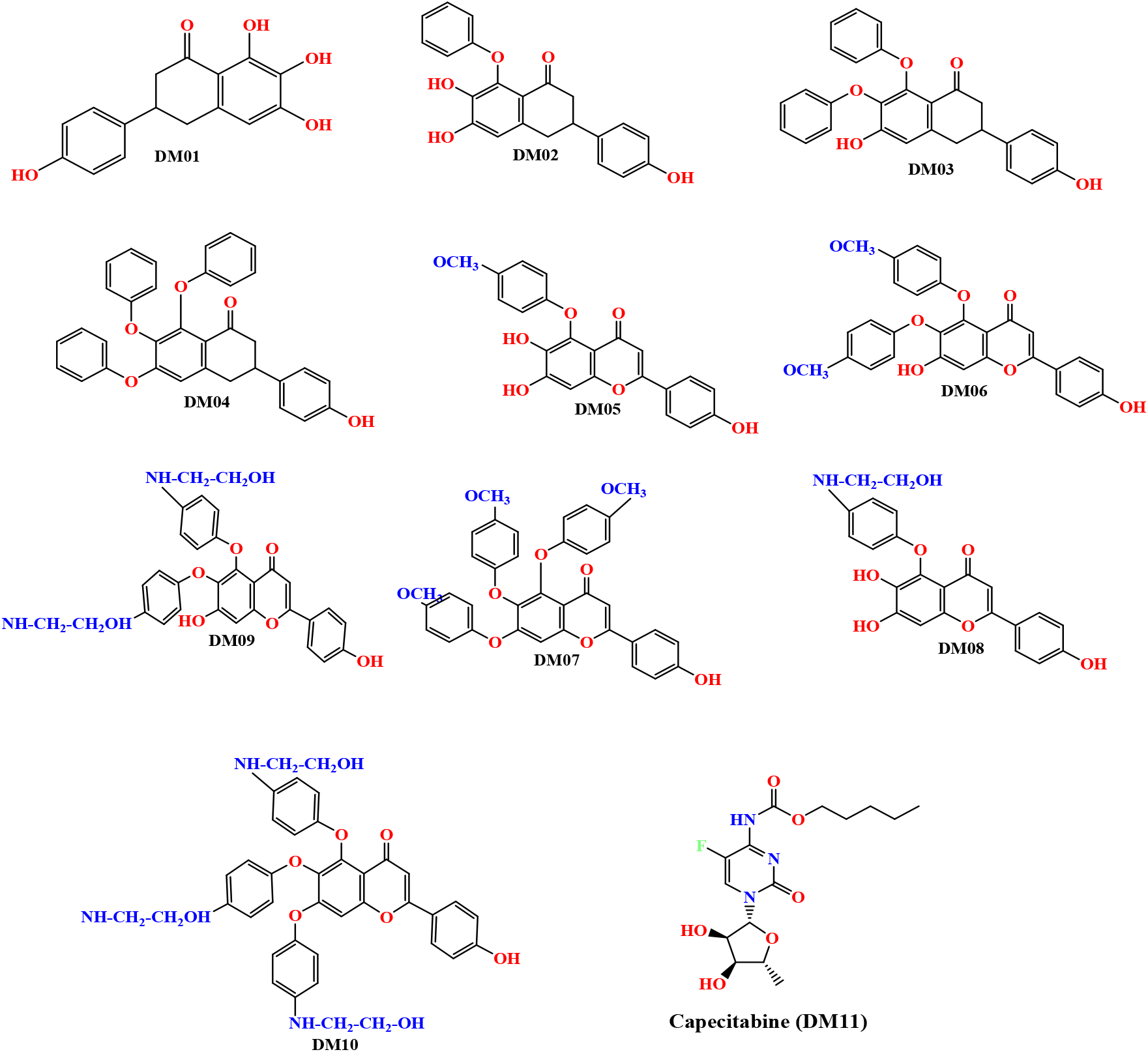
Chemical structure of Scutellarein derivatives

### 3.2 Optimized structure of selected scutellarin derivatives

The material studio application has been implemented to visualize the geometry optimization of the ten bioactive Scutellarein and its modification’s structure. The optimized chemical structures of these derivatives are displayed in (Fig.4). the optimized structure represents the symmetry and asymmetry point through the chemical structure and configuration. The goal of lead optimization is to maximize the efficiency of the most promising compounds to maintain the desired properties at optimum conditions [48].

**Figure 4.**
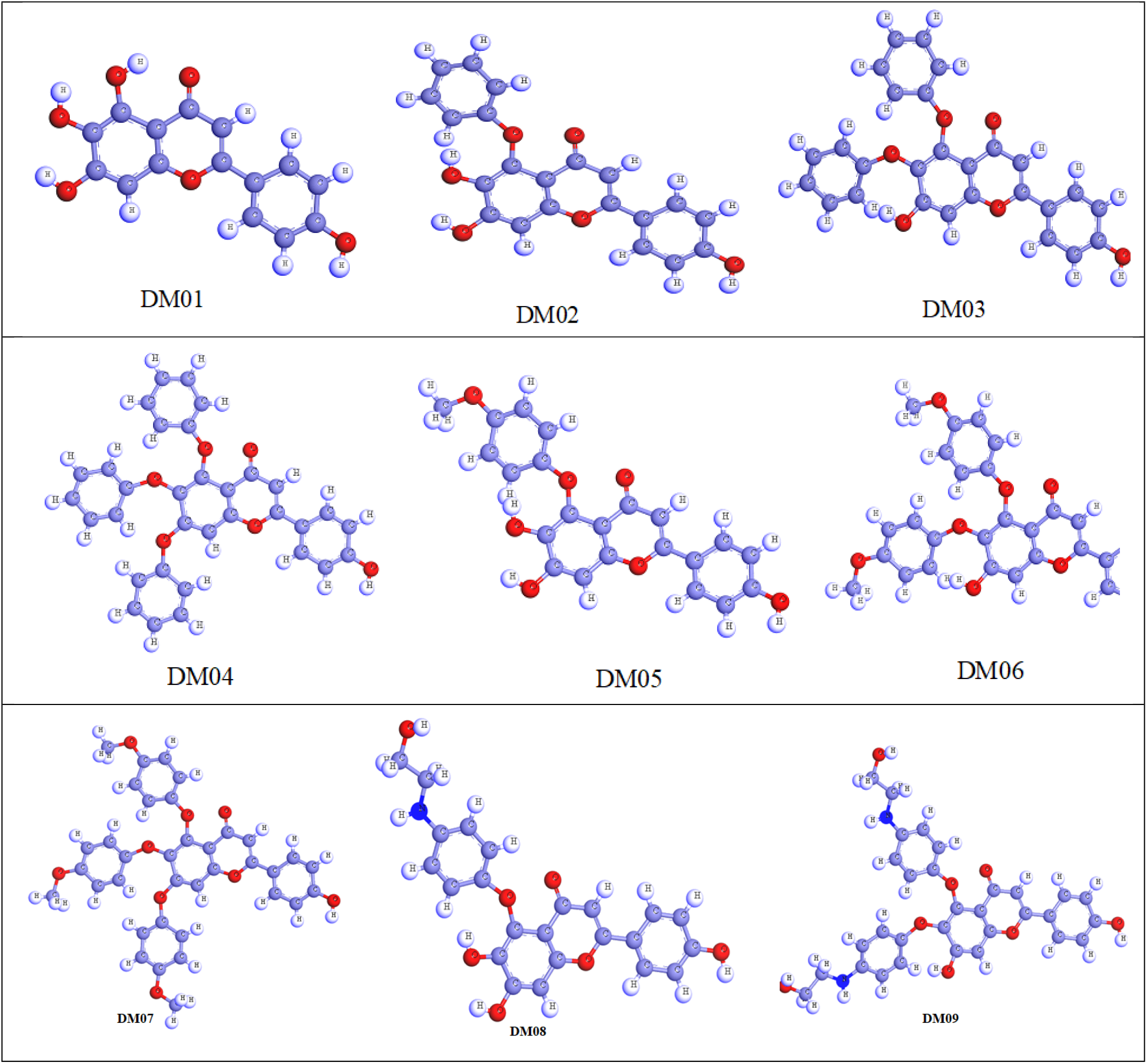
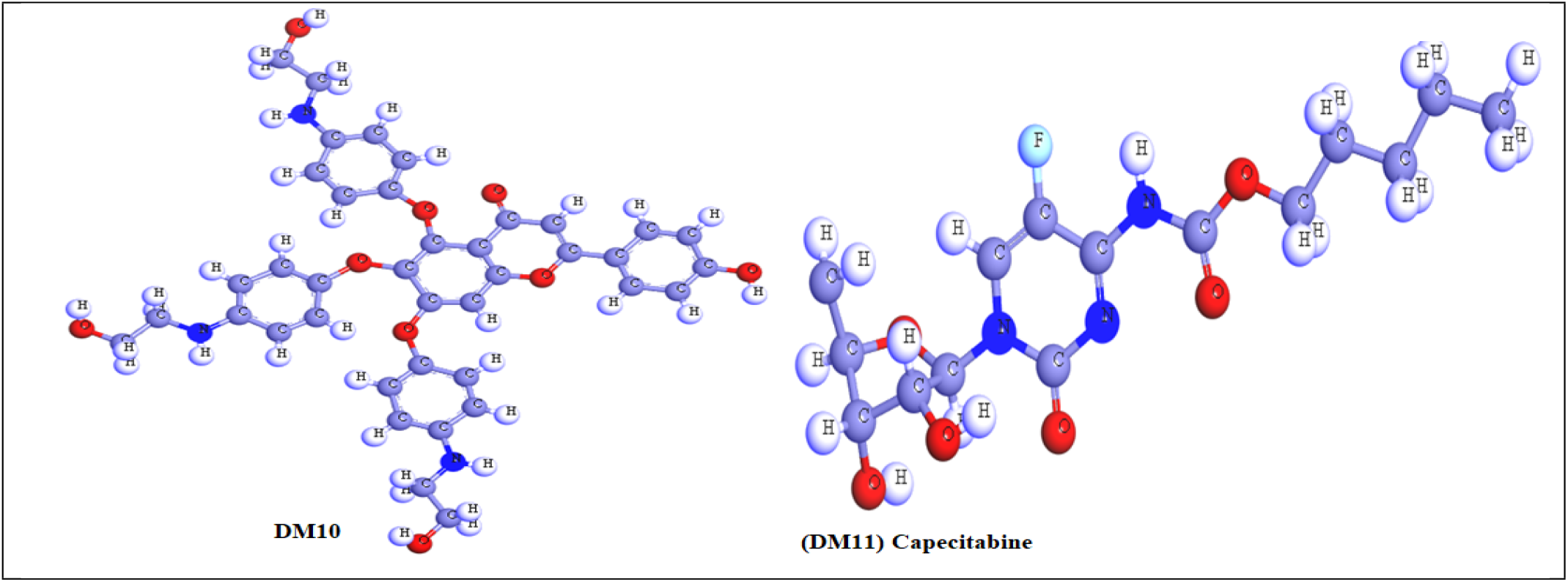
Optimized structure of Scutellarein derivatives

### 3.3 Pass Prediction spectrum

Pass is online software (which may be used to explore novel therapeutic actions for synthesized (materials(http://way2drug.com/PassOnline/predict.php)[49]. With the studied compounds, this implementation was carried out to detect possible therapeutic qualities that might be validated by laboratory investigation. PASS Prediction value evaluates the geometry of a novel molecule to the structures of well-known physiologically active substances, allowing researchers to determine whether or not a chemical will have a particular activity. This method can be applied from the very beginning of the investigation or development of new drugs. The previously generated geometries of the described molecules were uploaded to the PASS online tool as a mol form, and the potential mode of action and bioactivities were projected.

Regarding data, the probability of being active (Pa) value range is 0.161 to 0.383 for antiviral, 0.182 to 0.402 for antibacterial, 0.236 to 0.515 for antifungal, and the last one is antineoplastic, which is 0.572 to 0.794 (Showing in Table 2). It has been seen that the probability of being active (Pa) values are much more significant for antineoplastic than antiviral, antibacterial, and antifungal. So, after observing the data on the probability of being active (Pa), It has been decided to study them against cancer further. So, two proteins of triple-negative breast cancer proteins have been picked up as targeted receptors for further computational analysis.

**Table 1:** Effects of scutellarin on the proliferation of several cancer cell lines and the underlying molecular pathways[23].

| Cancer cell line | Molecular mechanism of scutellarin | Reference |
| --- | --- | --- |
| Colorectal cancer | Activation of caspase-6 resulted in the stimulation of cancer cell death and cancer cell spread. | [24] |
| Prostate cancer | Induced cell death (apoptosis) and induced cell cycle arrest in the G2/M phase to restrict cell growth. | [25] |
| Breast cancer | up-regulating the Hippo/YAP signaling pathway led to a suppression of cell proliferation and invading of the target cell. | [26] |
| Lung cancer | Elevated levels of apoptosis and autophagy in lung cancer cells are caused by activation of the ERK/p53 and c-met/AKT signal transduction pathways | [27] |
| Renal cell carcinoma | Suppressed renal cell carcinoma cell growth and invasion by stimulating phosphatase and tensin homologue expression. | [28] |

**Table 2:**
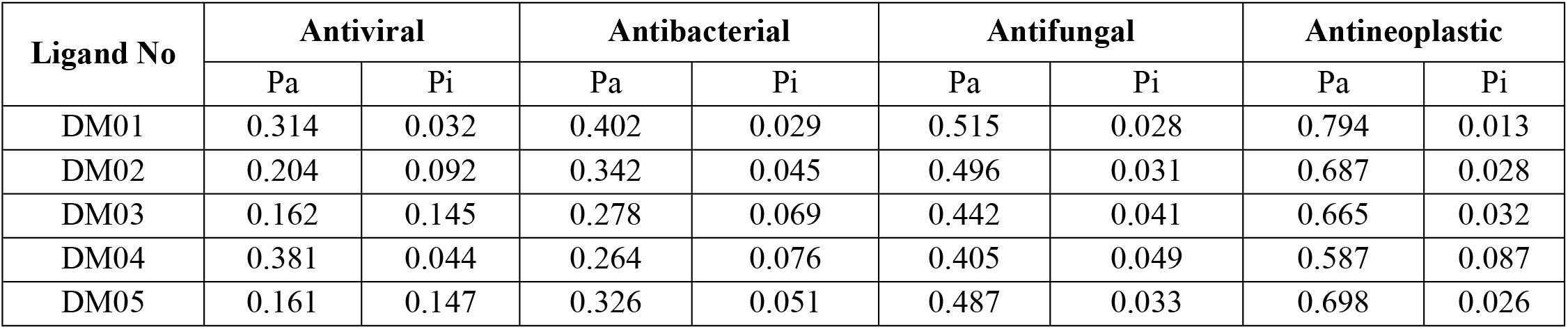
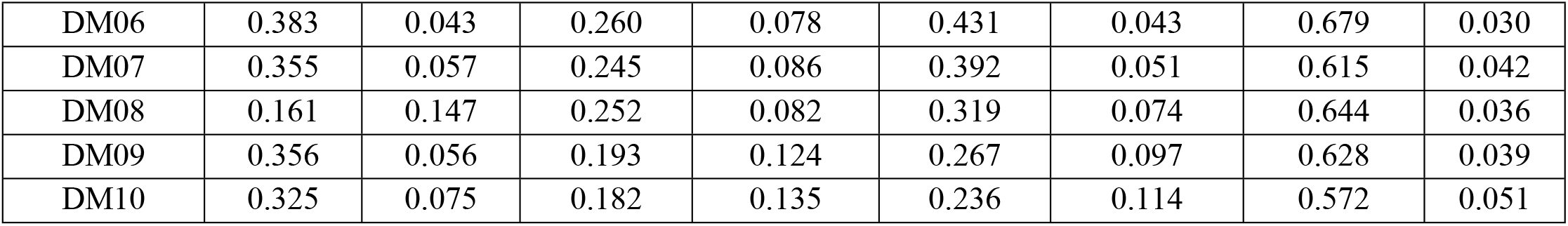
Computational data of pass prediction

### 3.4 Lipinski rule analysis for oral medication

Orally active drugs should be relatively small according to the Lipinski rule[50]. Lipinski’s rule of five establishes whether a biochemical molecule with a specified pharmacological or biological activity has molecular qualities and physical features, constituting it a probable orally administered medication in mammals[51]. The established Lipinski rule is following, Molecular weight should be less than 500, Octanol-water partition coefficient logP should be less than < 5 3., The number of hydrogen bond donors should be less than 5, and the number of hydrogen bond acceptors less than 10. The last one is Molar refractivity range should be 40 and 130.

According to this concept, medicine is considered to have suitable oral medication If they should fulfill this criterion. So, in our predicted data, it has been seen that the molecular weight of the compounds is below 286.24 to 691.73, and the range of topological polar surface area is 78.13 – 174.91. Besides, the bioavailability scores are better for all compounds (0.55); only ligands DM09 & DM10 are poor bioavailability scores. Another essential parameter is the GI absorption rate, which indicates that most drugs are poorly absorbed in the GI tract but the molecules DM01, DM02 & DM05 have high GI absorption rates (**Presented in Table 3**). Finally, it is observed that all drugs have satisfied the Lipinski rule where only two molecules declined by Lipinski rule DM09 & DM10 due to high molecular weight. So, ignoring the molecular weight, they should be recommended as potential oral medication.

**Table 3:** Predicted data of Lipinski rule

| Ligand No | Number of rotatable bonds | Hydrogen bond acceptor | Hydrogen bond donor | Topological polar surface area, Å<sup>2</sup> | Consensus Log Po/w | Lipinski rule |  | Molecular weight | Bioavailability Score | Gastrointestinal absorption |
| --- | --- | --- | --- | --- | --- | --- | --- | --- | --- | --- |
|  |  |  |  |  |  | Result | violation |  |  |  |
| DM01 | 01 | 06 | 04 | 111.13 | 1.81 | Yes | 00 | 286.24 | 0.55 | High |
| DM02 | 03 | 06 | 03 | 100.13 | 3.16 | Yes | 00 | 362.33 | 0.55 | High |
| DM03 | 05 | 06 | 02 | 89.13 | 4.68 | Yes | 00 | 438.43 | 0.55 | Low |
| DM04 | 07 | 06 | 02 | 78.13 | 6.20 | Yes | 01 | 514.52 | 0.55 | Low |
| DM05 | 04 | 07 | 03 | 109.36 | 3.17 | Yes | 00 | 392.36 | 0.55 | High |
| DM06 | 07 | 08 | 02 | 107.59 | 4.67 | Yes | 00 | 498.48 | 0.55 | Low |
| DM07 | 10 | 09 | 01 | 105.82 | 6.20 | Yes | 01 | 604.60 | 0.55 | Low |
| DM08 | 07 | 07 | 05 | 132.39 | 2.57 | Yes | 00 | 421.40 | 0.55 | Low |
| DM09 | 11 | 08 | 06 | 153.65 | 3.55 | No | 02 | 556.56 | 0.17 | Low |
| DM10 | 16 | 09 | 07 | 174.91 | 4.46 | No | 03 | 691.73 | 0.17 | Low |

**Table 4:** Data of chemical descriptors calculation.

| S/N | A=-LUMO | I=- HOMO | Energy Gap<br>$E(\text{gap}) = I - A$ | Chemical potential<br>$(\mu) = -\frac{I + A}{2}$ | Electronegativity<br>$(\chi) = \frac{I + A}{2}$ | Hardness<br>$(\eta) = \frac{I - A}{2}$ | Softness<br>$(\sigma) = \frac{1}{\eta}$ |
| --- | --- | --- | --- | --- | --- | --- | --- |
| DM01 | -1.645 | -8.648 | 7.003 | 5.1465 | -5.1465 | 3.5015 | 0.2856 |
| DM02 | -1.561 | -8.564 | 7.003 | 5.6250 | -5.0625 | 3.5015 | 0.2856 |
| DM03 | -1.579 | -8.205 | 6.626 | 4.8920 | -4.892 | 3.313 | 0.3018 |
| DM04 | -3.569 | -5.292 | 1.723 | 4.4305 | -4.4305 | 0.8615 | 1.1608 |
| DM05 | -3.422 | -6.651 | 3.209 | 5.0465 | -5.0465 | 1.6045 | 0.6231 |
| DM06 | -1.748 | -8.445 | 6.697 | 5.0965 | -5.0965 | 3.3485 | 0.2986 |
| DM07 | -1.674 | -8.426 | 6.752 | 5.0500 | -5.050 | 3.3760 | 0.2962 |
| DM08 | -1.572 | -7.494 | 5.922 | 4.5330 | -4.533 | 2.961 | 0.3377 |
| DM09 | -1.568 | -7.576 | 6.008 | 4.5720 | -4.5720 | 3.004 | 0.3329 |
| DM10 | -1.547 | -7.543 | 5.996 | 4.5450 | -4.5450 | 2.998 | 0.3336 |

### 3.5 Molecular Orbitals and Chemical Reactivity Descriptor

Chemical reactivity and kinetic stability are implied to be represented by frontier orbitals of a molecule, and it is the essential orbitals feature in molecules bioactivity. There are two types of frontier orbitals in molecules: HOMO and LUMO. The shift of the electron from the lowest to the highest energy state is mainly accounted for by the excitement of one electron from HOMO to LUMO[52]. Consequently, the excitement of electrons from the HOMO to the excited state LUMO implies significantly more energy. The kinetic stability of a molecule is a linear relationship between the HOMO-LUMO energy gap, which is described as increasing the HOMO-LUMO energy gap, simultaneously growing the chemical reactivity and kinetic stability [53].

Table 3 displays the computed values of molecular orbital energies, including the two well-recognized chemical variables of the energy gap, Chemical potential, Electronegativity hardness, and softness. The documented compound DM01 & DM02 implied the highest HOMO-LUMO energy gap (7.003 eV), which regards them as a more stable configuration. Besides, the derivatives with the maximum softness value were estimated to be -1.1608 in Ligand DM04, which means this drug could be dissolved more quickly. Simultaneously, the maximum hardness is about -3.5015 in Ligand DM01 & DM02, and this hardness data indicates that they might have required more time to break after reaching the physiological system. In addition, the stability of drugs, such as reactive species, molecules, and their propensity to molecularly react to create new substances, incorporate new physiochemical phenomena, and so on are all determined by their chemical potential under the most prevalent thermodynamic circumstance of constant temperature and pressure[53]. The calculated chemical potential ranges between 4.4305 and 5.6250, indicating a better score and a stable configuration has been displayed in **Table 4**.

Electronegativities are another vital parameter for drug design. In chemistry, polar bonds are a form of covalent bond. When the electronegativities of two or more atoms in a bond are very different from one another (>0.4), we indicate that the bond is polar. To put it another way, the negative charge of the electrons is not spread uniformly throughout the structure due to the nature of polar interactions[54]. In our investigation, the electronegativity ranges are obtained from - 4.4305 to -5.6250. So, it is stated that they should have uniformly distributed and shared the electron[55].

### 3.6 Frontier Molecular Orbitals (HOMO and LUMO)

The HOMO and LUMO orbitals have a crucial role in determining the efficiency of kinetics and having a biological impact on protein interactions.[56]. The HOMO and LUMO orbital geometries describe an electron-rich area and electron deficiency area. They were determined using the DFT method. **Fig. 5** shows that LUMO is often encountered in cation regions, notably in the positive atom-containing compartments where the positive charge has been located. Thus, an additional electron is attached to the LUMO segment. On the other hand, HOMO is often encountered in anion segments, notably in the negative atom-containing compartments where the electronegative charge could be presented. To illustrate, the yellow and deep maroon moiety displayed the LUMO or positive charge region, and the bright greenish shade revealed the HOMO segment or negative charge portion. It is often thought that the protein may be coupled to the LUMO portion of the molecules[57, 58].

**Figure 5.**
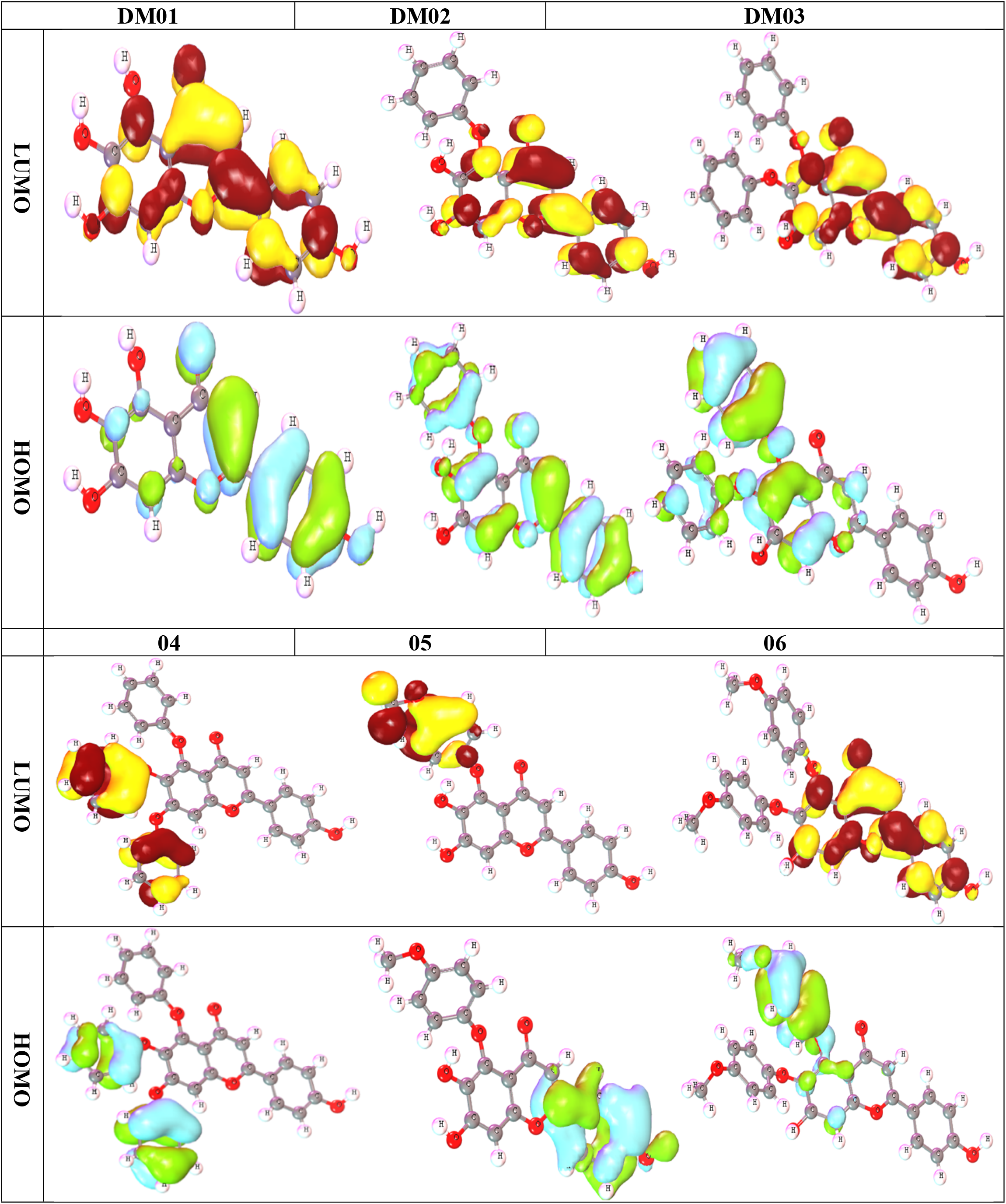
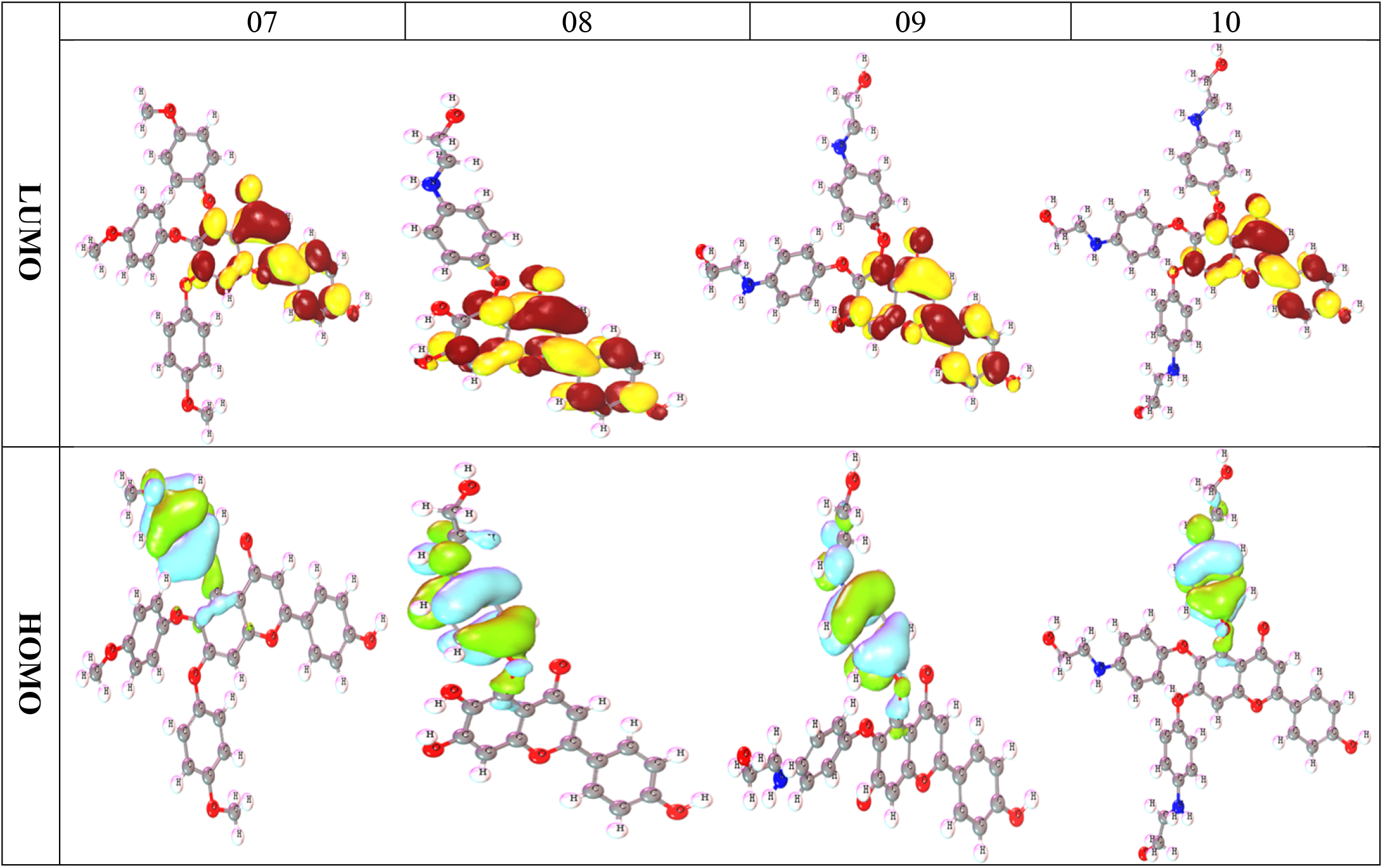
Frontier Molecular Orbitals (HOMO and LUMO) diagram

### 3.7 Molecular of Electrostatic Potential (MEP) charge distribution mapping

The MEP has gained a compatibility aspect that shows the most promising zones for the targeted electrophile and nucleophile of charged area molecules on organic materials[59]. The MEP clarifies the biological mechanism and the H-bonding coupling or interaction[60] and it is significant in the investigation of describing organic molecules. The electrostatic potential map represents a simple approach for determining the interactions between different molecular geometries. In this study, the electrostatic potential map of the targeted compound has been simulated using B3LYP, with a basis set of 3-21G, and optimized the structure as an outcome. (Fig. 6). It displays the molecular structure and size and positive, negative, and neutral electrostatic probability regions by displaying the color difference. Besides, it is also a prominent approach to investigating the relationship between physicochemical characteristics and the structure of the targeted compound[61, 62].

**Figure 6.**
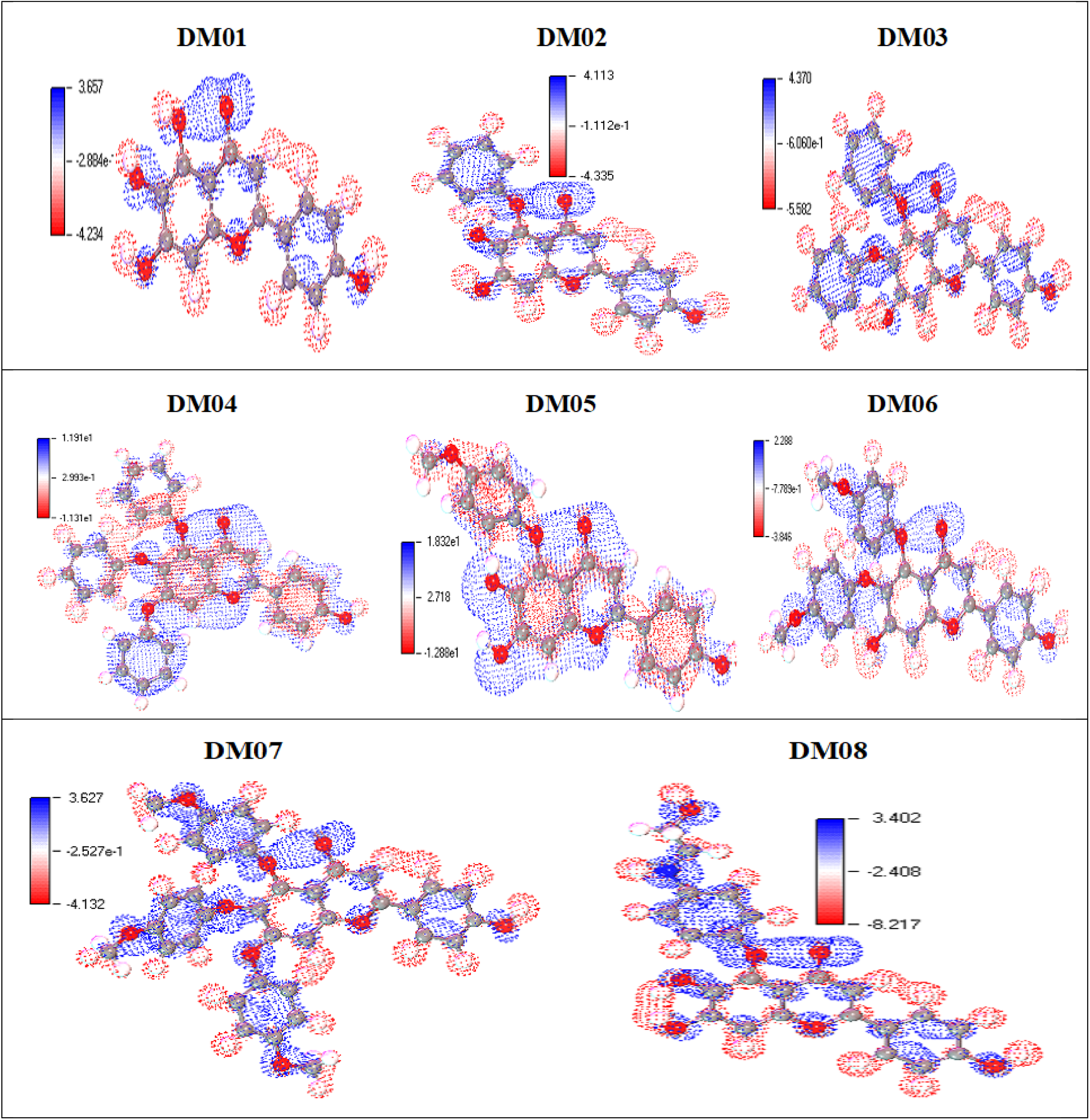
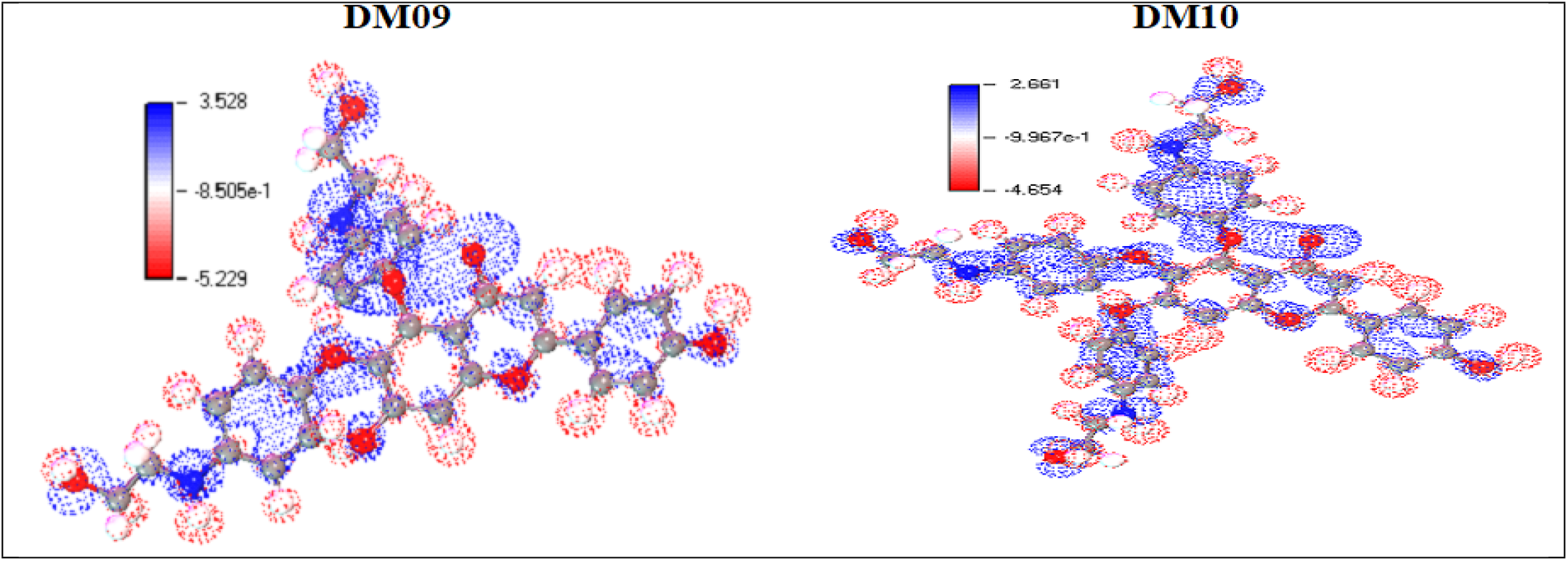
Molecular electrostatic potential (MEP) mappings

The assaulting region’s potential declines in the following manner: blue, red, and white. The white region is indicated as the neutral region where no assault occurs. The red color shows the high electron saturation area, and this zone described that electrophile might readily assault. The minimal electron density surface is represented by the blue hue, which is amenable to nucleophilic assault.

### 3.8. Molecular docking against Triple-negative Breast cancer

The docking computations were conducted with the help of the program AutoDock (PyRx), and the binding energy of the peptide was determined. For the interactions between drugs and receptors have been listed in **Table 5**. The standard binding affinity has been considered an active drug molecule of -6.0kcal/mol [62-66]. The Scutellarein derivatives (DM03, DM04, DM06, and DM07) were found to have the maximum bioactivity against triple negative Breast cancer (PDB ID 7L1X) & compounds (DM02. DM05 & DM08) have shown the most potency when compared to the all-other derivatives.

**Table 5:** Data of binding energy against triple-negative breast cancer.

| Drug Molecules No | H Human CK2 alpha kinase (PDB ID 7L1X) |  |  | Triple-negative Breast cancer receptor (PDB ID 5HA9) |  |  |
| --- | --- | --- | --- | --- | --- | --- |
|  | Binding Affinity (kcal/mol) | No of H Bond | No of Hydrophobic Bond | Binding Affinity (kcal/mol) | No of H Bond | No of Hydrophobic Bond |
| DM01 | -9.5 | 04 | 11 | -9.4 | 02 | 05 |
| DM02 | -9.7 | 01 | 11 | -10.5 | 04 | 04 |
| DM03 | -10.7 | 01 | 11 | -8.2 | 03 | 06 |
| DM04 | -11.0 | 02 | 07 | -7.6 | 02 | 06 |
| DM05 | -9.6 | 04 | 11 | -10.5 | 02 | 06 |
| DM06 | -10.0 | 02 | 12 | -7.7 | 02 | 09 |
| DM07 | -10.0 | 04 | 17 | -8.1 | 02 | 09 |
| DM08 | -9.2 | 03 | 12 | -10.0 | 06 | 04 |
| DM09 | -9.1 | 06 | 11 | -7.5 | 05 | 07 |
| DM10 | -8.8 | 06 | 12 | -7.6 | 08 | 04 |
| Standard<br>Capecitabine | -7.7 | 03 | 03 | -7.5 | 02 | 06 |

For triple-negative breast cancer (PDB ID 7L1X), the most significant binding affinity has been seen at 11.0 kcal/mol, and -10.7 kcal/mol in ligand DM03, and DM04, and the most considerable binding affinity has been seen at -10.5 against (PDB ID 5HA9) in ligand DM02. So, to compare with standard drugs, capecitabine has also been studied with the synthetic ligand. It is found that the new drug molecules are more active than the FDA-approved drugs capecitabine. So, it can be said that the drug molecules are opposed to the standard binding affinity, actively bind with triple-negative breast cancer protein as a potential inhibitor, and can be used as an effective drug.

### 3.10 Protein-ligand interaction

The interaction diagrams of the drug-protein combinations, hydrogen bonding, and molecular docking pocket have been developed utilizing software BIOVIA Discovery Studio and Pymol application. The hydrogen bond interactions such as conventional and non-conventional H bonds, hydrophobic interactions including pi-sigma, alkyl, and pi-alkyl interactions, and hydrogen bond donor and hydrogen bond -acceptor interactions have been generally looked at protein and ligand interaction. Different engagement and binding energies between the substance and its desired targeted protein are represented graphically in **Fig. 7**.

**Figure 7.**
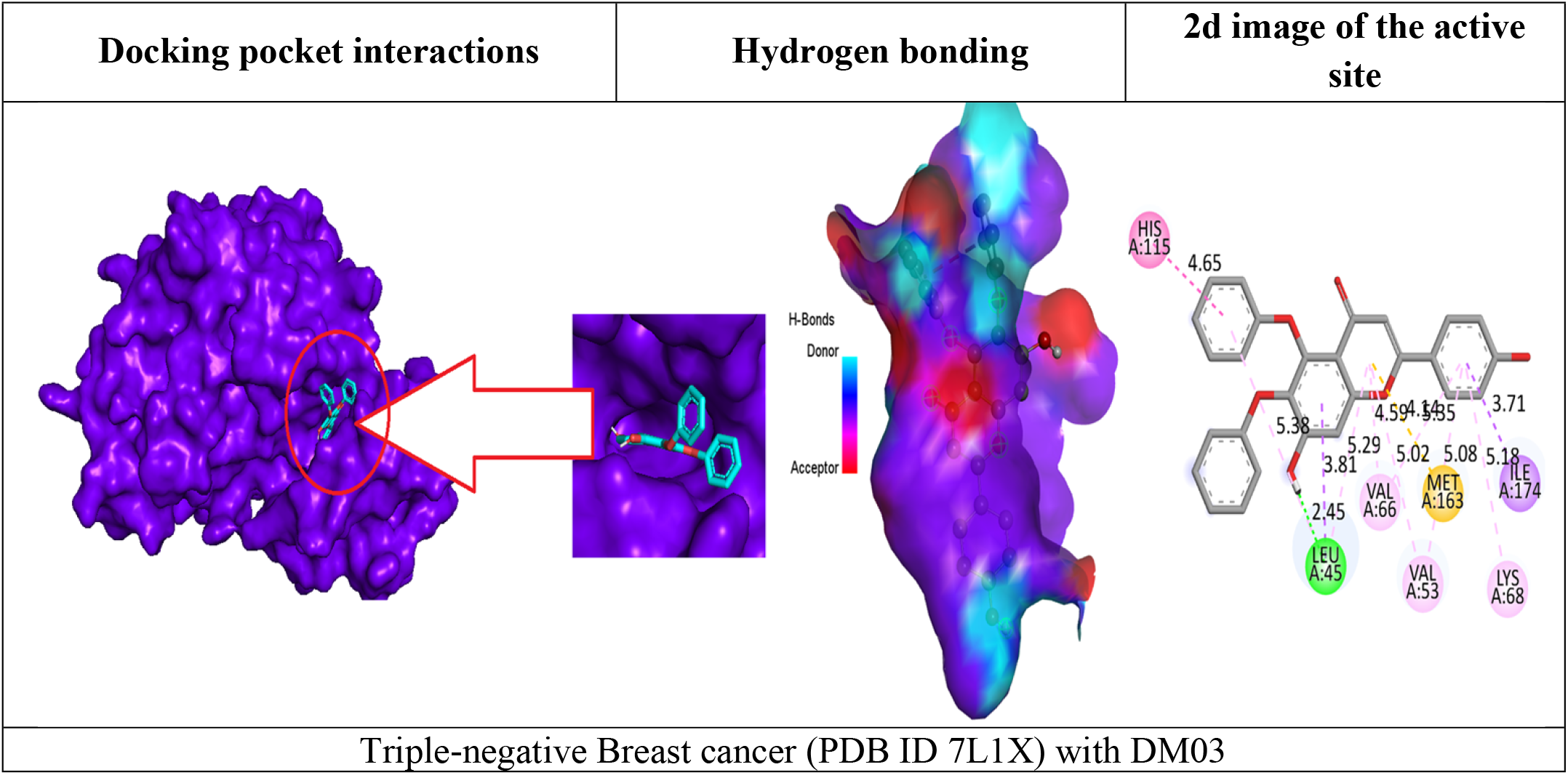
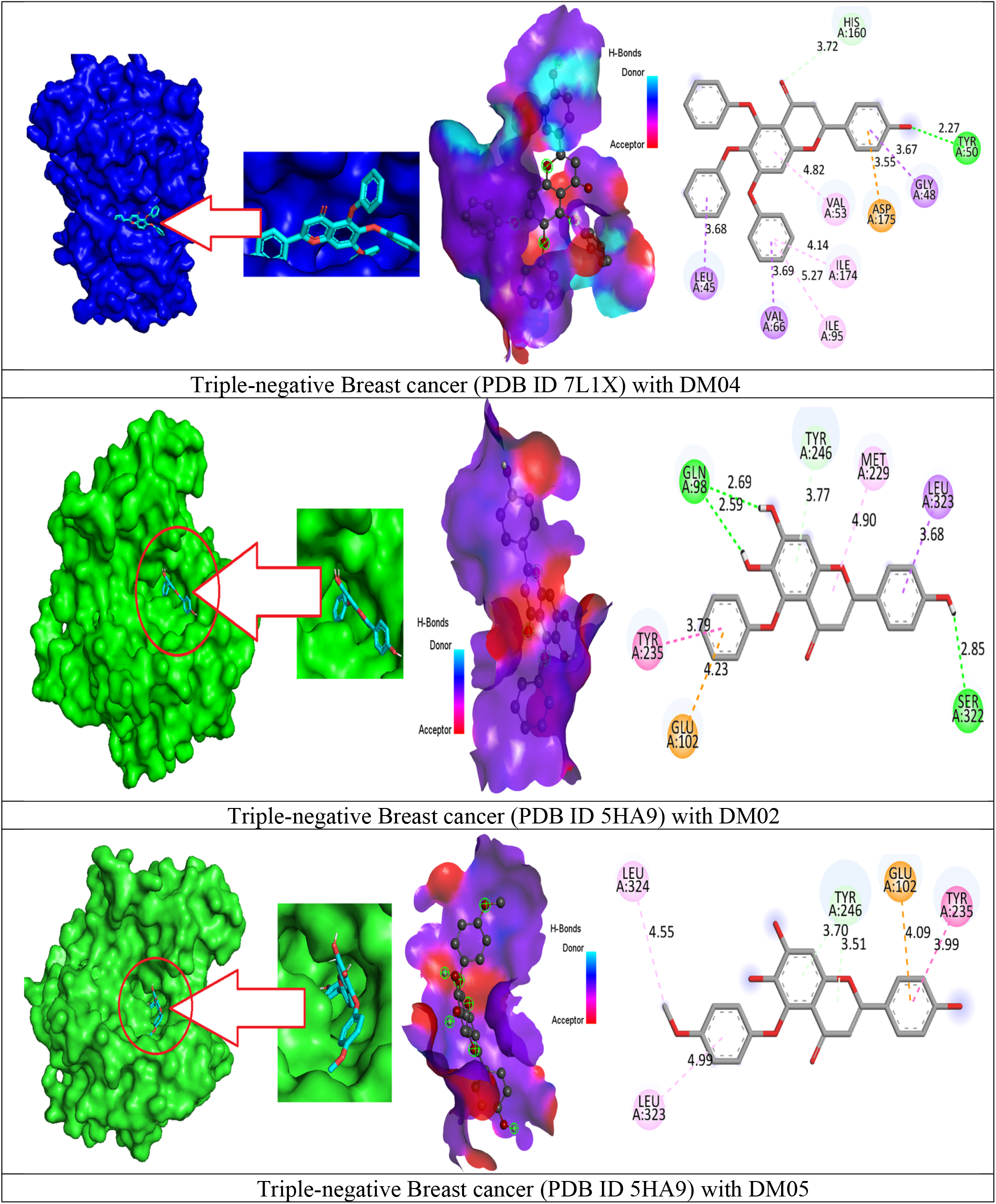
The results of Molecular docking pose interactions between compounds and protein.

In hydrogen bonding, the red color is described acceptor region, whereases the acceptor region describes the sky-blue color. Besides, the 2d image of active amino acid residues is seen that A: TYR-50, A: HIS-160, A: ASP-175, A: LEU-45, A: GLY-48, A: VAL-66, A: VAL-53, A: ILE-95, and A: ILE-174 is generated for triple negative Breast cancer (PDB ID 7L1X) with DM04, whereases A: TYR-50, A: HIS-160, A: ASP-17,5 A: LEU-45, A: GLY-48 A: VAL-66, A: VAL-53, A: ILE95, and A: ILE-174 is generated during the formation of the complex with PDB ID 5HA9. Besides, the amino acid residues and their bond distance for all compounds are given in the supplementary file.

### 3.11 Molecular Dynamic simulation

Molecular dynamics have been simulated protein-ligand complexes to express high stabilization at the simulation point of the compounds and determine the accuracy docking procedure in terms of the average root-mean-square deviation (RMSD) and root-mean-square fluctuation (RMSF). This value represents the binding pose in the respective crystal structures, ligand, and protein interaction complex. The RMSD and RMSF drift assessment statements have been implemented to determine the molecule’s stability. **Fig. 8** illustrate the diagram. The MD simulation point indicates that the drug complex has excellent engagement in the drug potential pocket.

**Figure 8.**
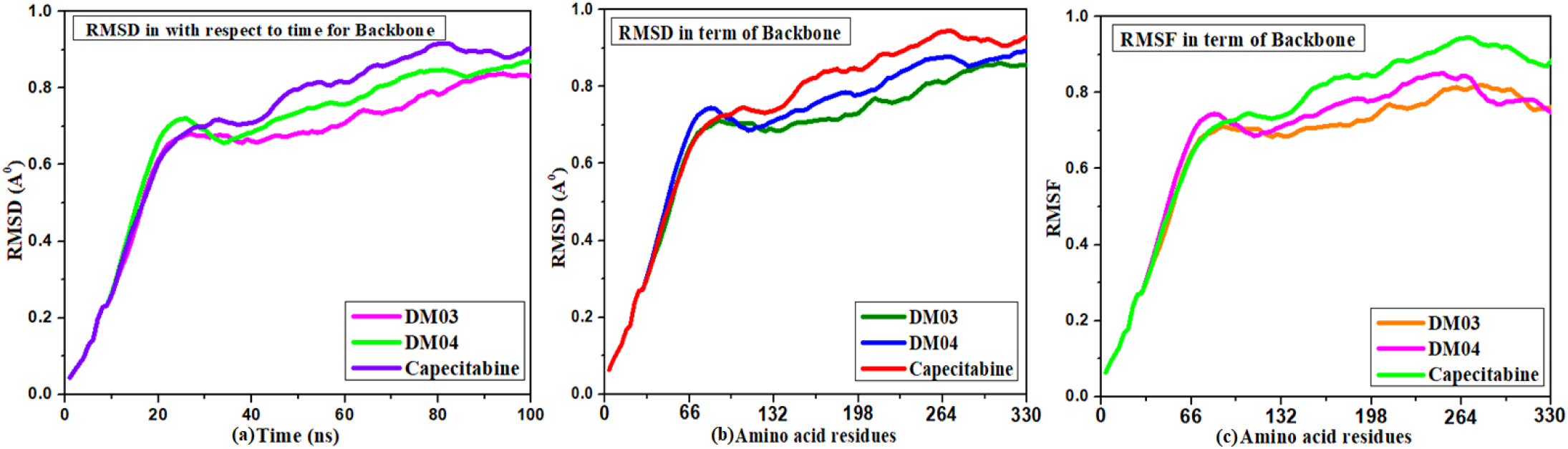
MD simulation and RMSD, RMSF results of triple negative Breast cancer with ligand.

For triple negative Breast cancer (PDB ID 7L1X), the RMSD scatterplot in **Fig.8(a)** demonstrates that the dynamic evaluation of picked molecules from the beginning to the end position is consistent from 0.8 Å to 1.0 Å throughout the simulation period. RMSD of docked complexes has been run at 100ns. Again, the RMSD in terms of amino acid backbone **Fig.8 (b)**, 0.9 Å, and RMSF is 0.8 Å as maximum **Fig.8 (c)**. It has been observed that the range of MD simulation is much lower for the newly designed compounds DM03 & DM04 compared to the standard drug capecitabine. This value indicates the reported drug is highly stable.

### 3.12 ADMET data prediction

ADME-related features such as therapeutic absorption, distribution, metabolism, solubility, and even oral bioavailability are influenced by ADMET characteristics. The computational inspection approaches might be used in the drug development process to predict ADMET features. Water solubility is the most potent criterion for oral drug delivery systems in modern drug discovery. High water-soluble oral drugs consistently have excellent oral bioavailability with maximum absorption capability[67]. In order to estimate solubility in water for active drug candidates, there is a great interest in developing quick, reliable, structure-based approaches.

In the listed ADMET feature, the water solubility of each compound is a different value. The ligands DM04, DM07, DM08 & DM10, are highly soluble in water, and their range is from -2.893 to -2.998 since the actual values of water solubility (Log S) for slightly soluble vary from -4 to -6, and high solubility compounds -2 to -4, correspondingly[68]. The remaining compounds are little soluble in water which means they are highly soluble in fatty material or lipid. Another absorption parameter is Caco-2 permeability. The standard range of good Caco-2 Permeability is >0.90 according to the pkCSM web tool[69]. In our finding, all the drug has an excellent absorption rate in Caco-2 Permeability, excluding DM04 since it has been higher Caco-2 Permeability than 0.9.

**Table 6** lists the distributional features of all substances, including their volume distribution and BBB permeability. The volume distribution of medicine is closely related to medication distribution. The volume of distribution (VD) of a drug throughout blood and tissue should be used to estimate whether the drug distributes uniformly or not uniformly. Lower VD indicates that the drug concentration in plasma is more significant, and the medication cannot diffuse to the tissues as effectively [70]. In this case, most molecules had low VD levels, while some had intermediate VD values. Another parameter, the blood-brain barrier (BBB), is a protective barrier that inhibits unwanted materials from entering the brain and central nervous system (CNS)[71].In our finding, no drug molecules can permeability BBB.

**Table 6:** ADMET profiling prediction of the studied ligands

| Molecules No | Absorption |  | Distribution |  | Metabolism |  | Excretion |  |
| --- | --- | --- | --- | --- | --- | --- | --- | --- |
|  | Water<br>solubility<br>Log S | Caco-2<br>Permeability<br>10<sup>-6</sup> cm/s | VDs (human) | BBB<br>permeability | CYP450 1A2<br>Inhibitor | CYP450 2C9<br>substrate | Total<br>clearance<br>(ml/min/kg) | Renal OCT2<br>substrate |
| DM01 | -3.064 | -0.004 | -0.193 | No | Yes | No | 0.511 | No |
| DM02 | -4.235 | 0.416 | -1.10 | No | Yes | Yes | 0.311 | No |
| DM03 | -3.443 | 0.915 | -1.267 | No | No | Yes | 0.385 | No |
| DM04 | -2.977 | 0.990 | -0.87 | No | No | Yes | 0.466 | No |
| DM05 | -3.087 | 0.85 | -0.584 | No | Yes | Yes | 0.718 | No |
| DM06 | -3.101 | 0.224 | -1.307 | No | No | Yes | 0.792 | No |
| DM07 | -2.893 | -0.159 | -0.637 | No | No | Yes | 0.973 | No |
| DM08 | -2.998 | 0.677 | -0.388 | No | Yes | Yes | 0.668 | No |
| DM09 | -3.029 | 0.26 | -1.301 | No | No | Yes | 0.664 | No |
| DM10 | -2.896 | -0.472 | -0.851 | No | No | No | 0.643 | No |
| Capecitabine | -2.939 | 0.339 | -0.253 | No | No | No | 1.138 | No |
| Fluconazole | -2.66 | 1.104 | -0.624 | No | Yes | No | 0.433 | No |

On the other hand, all the ligands may actively metabolize in CYP450 2C9 substrate while only DM01, DM02, DM05 & DM08 are Inhibited by CYP450 1A2.Finally, the total clearance range is about 0.220 (ml/min/kg) to 0.792 (ml/min/kg) which mean maximum 0.792 (ml/min/kg) drug can out from the body and no drug can excrete through Renal OCT2 substrate. This computational ADMET projection research suggests that the new molecules DM01–DM10 are biologically better drug candidates, which should be considered promising as new drug molecules.

### 3.13. Aquatic and non-aquatic toxicity

Another parameter is aquatic and non-aquatic toxicity. Aquatic and non-aquatic toxicity is a significant objective in assessing undesirable impacts on the environment or human use [72]. In this investigation, a series of computational methods for evaluating biochemical aquatic and non-aquatic toxicity have been discussed with different parameters such as AMES toxicity, oral rat acute toxicity (LD50), oral rat chronic toxicity, etc. In this case, it has been proven that all the drugs are free from AMES toxicity without DM02. The maximum tolerated dose for a human is 0.781 mg/kg/day in DM01. At the same time, the lowest maximum tolerated dose is 0.358 mg/kg/day in Ligand DM09, which indicates that the highest dose of 0.781 mg/kg/day should be taken for a given period (24 Hours); otherwise, the adverse effect may produce. According to current estimates, the acute oral toxicity and the oral rat chronic toxicity of the active compounds vary, with values ranging from 2.418 mg/kg/day to 3.43 mg/kg/day being documented for the oral rat acute toxicity, where the oral rat chronic toxicity being reported from -0.086 mg/kg/day to 4.029 mg/kg/day. Finally, they all are free from skin sensitization with the lowest T. pyriformis toxicity. So, it can be concluded that they might have no adverse effect on aquatic and non-aquatic and could be performed further experimental works in laboratories. Aquatic and non-aquatic poisoning are all of the medications listed in **Table 7**.

**Table 7:**
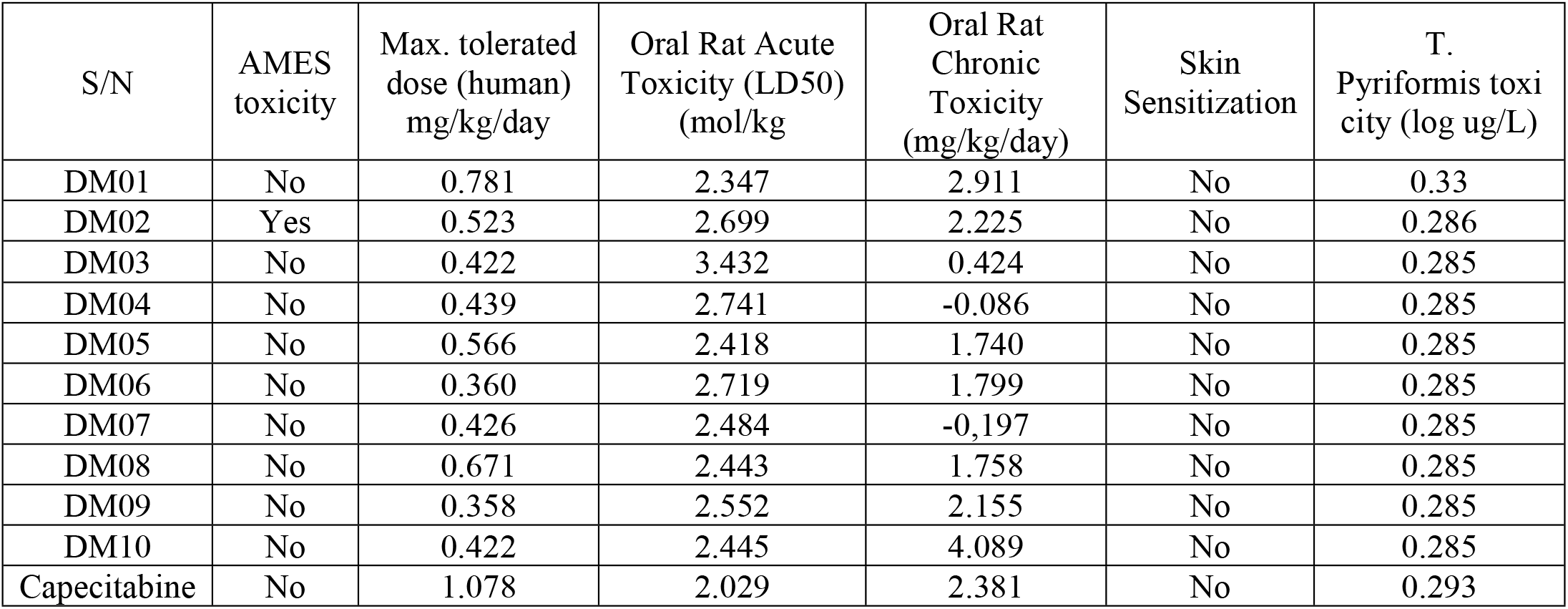
Summary of the aquatic and non-aquatic toxicity properties of the studied ligands

## 4. Conclusion

The study provides a computational examination of the pharmacological effects of 10 scutellarin derivatives against triple-negative breast cancer. The results of the cancer-fighting tests suggested that the newest scutellarin derivatives would be effective against triple-negative breast cancer in various ways. As mentioned earlier, the PASS findings we acquired for our medications demonstrated stronger potency against antineoplastic than viruses, fungi, and bacteria. It was decided to focus primarily on triple-negative breast cancer. For triple-negative breast cancer (PDB ID 7L1X), the docked compound of produced molecules demonstrates the highest binding affinities and excellent non-bonding (DM03 -10.7 kcal/mol, DM04-11.0 kcal/mol, DM06-10.0 kcal/mol, & DM07 -10.0 kcal/mol). Molecular electrostatic potential, HOMO, LUMO, the HOMO-LUMO gap, and other chemical reactivity features have been fine-tuned for biological use. The medications’ softness values ranged from -1.6045 to -2.2856, indicating that they would be metabolized rapidly once they entered the body. Due to their smaller HOMO-LUMO energy disparities, modified scutellarin derivatives like capecitabine will be more reactive than traditional ligands. This study’s crucial and pivotal aspects of the ADME and toxicity data. Results from the ADMET study showed that the novel compounds are safer than the reference medications concerning toxicity, water solubility, distribution volume, and clearance rate and that they do not have any carcinogenic effects due to the absence of DM02. Improved scutellarin ADMET characteristics compared favorably to gold-standard medicines. It is possible, finally, that these scutellarin derivatives are a valuable and viable treatment candidate against triple-negative breast cancer. Scutellarin should be used in more experiments to produce synthetic drugs to combat lethal TNBC.

## Funding

No funding was received in this study.

## Data Availability

All data are available in the text

## Conflicts of Interest

The authors declare that the research was conducted in the absence of any commercial or financial relationships that could be construed as a potential conflict of interest.

## Acknowledgement, and Funding statement

Author would like to acknowledge Department of Pharmacy, Faculty of Allied Health Sciences, Daffodil International University, Ashulia, Dhaka, Bangladesh, where the principal work has been being carried out. (Research4Life Group A Country: Bangladesh).

## Consent for Publication

No human or animal testing is referred; therefore, there is no need for ethical approval or informed consent.

## Notes

### Competing Interest Statement

The authors have declared no competing interest.

